# Loss of MAX1 redirects, rather than delays, the leaf senescence program in lettuce

**DOI:** 10.64898/2026.08.27.747486

**Authors:** Adi Kedem, Guy Azrieli, Mily Ron, Noam Ozeri, Megan Reeves, Dor Russ, Richard Michelmore, Lior Tal

## Abstract

**Background:** Strigolactones (SLs) regulate diverse aspects of plant development and have been implicated in promoting leaf senescence. However, senescence phenotypes associated with SL deficiency have not been consistently observed across species, suggesting that this function may be species- or context-dependent. Moreover, the contribution of endogenous SL biosynthesis to senescence in leafy vegetable crops remains unclear. Here, we investigated the role of the SL biosynthetic gene MORE AXILLARY GROWTH1 (MAX1) in dark-induced leaf senescence in lettuce (*Lactuca sativa*).

**Results:** We found that endogenous SL biosynthesis plays a major role in dark-induced senescence in lettuce. SL pathway genes were induced during dark storage, while exogenous GR24 accelerated senescence and lettuce MAX1 (LsMAX1) complemented the delayed-senescence phenotype of the *Arabidopsis max1* mutant. Consistent with these findings, CRISPR/Cas9-generated *Lsmax1* mutants exhibited a pronounced stay-green phenotype during prolonged darkness, accompanied by strongly reduced induction of key senescence-associated genes. Despite this delayed visible senescence, *Lsmax1* retained a substantial transcriptional response to dark storage. Strikingly, loss of LsMAX1 did not simply weaken the wild-type senescence program, but redirected part of the response toward a distinct stress-associated transcriptional state that was largely absent from wild type. Loss of LsMAX1 did not affect vegetative rosette architecture, although increased branching emerged after bolting.

**Conclusions:** Our findings establish MAX1-dependent SL biosynthesis as an important regulator of leaf senescence in lettuce and reveal a role that extends beyond controlling the rate of senescence. Rather than simply delaying the wild-type program, loss of LsMAX1 alters the transcriptional trajectory of senescence, favoring an alternative stress-associated state during prolonged darkness. The strong stay-green phenotype without detectable changes to vegetative rosette architecture further highlights SL biosynthesis as a potential target for extending postharvest longevity in lettuce and other leafy crops.

## Introduction

Strigolactones (SLs), carotenoid-derived terpenoid lactones, have emerged as positive regulators of leaf senescence across multiple plant species. In Arabidopsis, rice, and bamboo, genetic and pharmacological evidence consistently demonstrates that SLs promote dark-induced chlorophyll degradation and accelerate senescence progression [1–4]. Forward genetic studies identified ORE9 as MORE AXILLARY GROWTH 2 (MAX2), directly linking SL signaling to leaf longevity [5–7]. Exogenous application of the synthetic SL analog GR24 restores or accelerates senescence in SL-deficient backgrounds in Arabidopsis and rice [1, 2], and mutation of an SL biosynthesis gene in petunia delays leaf senescence [8]. Yet the generality of this function remains unresolved: clear senescence phenotypes have not been reported in SL mutants of pea or tomato [9–13], suggesting that the SL-senescence axis may be species- or context-dependent. Critically, whether endogenous SL biosynthesis contributes to leaf senescence in any leafy crop species has never been directly tested.

SL biosynthesis begins with the conversion of beta-carotene to carlactone through the sequential action of DWARF27 (D27) and the carotenoid cleavage dioxygenases CCD7/MAX3 and CCD8/MAX4 [14–17]. Carlactone is subsequently oxidized by the cytochrome P450 monooxygenase MORE AXILLARY GROWTH1 (MAX1) to form carlactonoic acid [15, 18], a central branch point in SL biosynthesis. In Arabidopsis, carlactonoic acid can be methylated by CARLACTONOIC ACID METHYLTRANSFERASE (CLAMT) to produce methyl carlactonoate, which is further modified by LATERAL BRANCHING OXIDOREDUCTASE (LBO) to generate bioactive noncanonical SL derivatives [19, 20]. In parallel, members of the CYP722 family catalyze alternative oxidations of carlactonoic acid, leading to the production of canonical SLs in many angiosperms [21]. These diversification steps all occur downstream of MAX1, positioning it at a metabolic pivot that determines flux toward structurally and functionally distinct SL end products.

SL perception and signal transduction are mediated by the α/β hydrolase receptor DWARF14 (D14), which binds SL and promotes association with the F-box protein MAX2 [22–25]. This interaction assembles an SCF-type E3 ubiquitin ligase complex that targets SMXL/D53 transcriptional repressors for ubiquitin-mediated degradation [26–30], thereby activating downstream transcriptional programs associated with SL responses, including senescence [31]. The stay-green phenotypes of *max2* and *d14* mutants confirm that intact SL perception is required for normal senescence progression [1]. Whether this requirement reflects control over the rate of senescence, or over which senescence-associated genes are engaged, has not been resolved.

SL levels are strongly influenced by nutrient and carbon availability. Phosphate deficiency consistently induces SL biosynthetic gene expression and increases SL accumulation across species [32–36], and SL signaling contributes to phosphate starvation transcriptional reprogramming in Arabidopsis [34]. SL biosynthesis is similarly linked to carbon status. During extended darkness, sugar starvation markers are induced prior to the upregulation of SL biosynthetic genes such as *MAX1* and *MAX3* [37], placing SL induction temporally downstream of carbon depletion, and exogenous sugars suppress SL-induced senescence under dark conditions [4]. Together, these findings position SL as an integrator of nutrient and carbon status, and suggest that the sustained darkness and carbon depletion characteristic of postharvest storage may represent conditions under which endogenous SL biosynthesis is engaged.

SL signaling further reinforces senescence-associated transcriptional programs in concert with ethylene and salicylic acid [1, 38]. These studies have primarily characterized SL action through its crosstalk with other hormone pathways, leaving open whether SL signaling also shapes the transcriptional architecture of senescence itself.

Despite this mechanistic understanding in model systems, harnessing SL biosynthesis for crop improvement has been constrained by architectural penalties. In several crop species, disruption of SL biosynthesis, signaling or transport results in excessive tillering or branching, including rice [33], tomato [11, 21, 39], and pea [9]. In tomato, SL biosynthesis mutants display severe shoot overbranching accompanied by reduced yield, and restoration of agronomic performance required grafting a wild-type shoot onto the mutant rootstock [39]. These findings illustrate the challenge of exploiting SL pathway manipulation for crop improvement without incurring undesirable architectural trade-offs. Whether SL-dependent suppression of axillary branching is conserved in lettuce, and critically, whether it is temporally restricted to specific developmental stages, has not previously been reported. Lettuce offers a distinct opportunity to address both questions: it is harvested at the vegetative rosette stage prior to bolting, and its commercial value resides entirely in leaf quality rather than reproductive architecture, allowing the direct contribution of SL biosynthesis to leaf senescence to be examined in a crop context for the first time.

Postharvest senescence in lettuce is driven by sustained darkness and carbon limitation following harvest, leading to rapid chlorophyll degradation and metabolic decline that restrict shelf life. Cytokinin-based strategies such as IPT expression have been used to delay senescence [40]. Exogenous application of the synthetic strigolactone analog GR24 was recently reported to improve postharvest quality and delay chlorophyll loss in celery [41]. However, the role of endogenous strigolactone biosynthesis in postharvest leaf longevity of leafy vegetables has not been directly tested genetically. Here, we investigate whether *LsMAX1*-dependent SL biosynthesis contributes to dark storage-induced senescence in lettuce by characterizing a CRISPR/Cas9-generated loss-of-function mutant in *LsMAX1*. We further use RNA sequencing to characterize how loss of *LsMAX1* redirects the leaf senescence transcriptional program.

## Results

### SL pathway genes are induced during postharvest storage and GR24 accelerates senescence in lettuce

To assess whether SL signaling is engaged during postharvest senescence in lettuce, we profiled the senescence marker *LsORE1* alongside core SL pathway genes across the storage time course by RT-qPCR. *LsORE1* was strongly induced during storage, confirming that senescence progressed over this time course. The SL biosynthetic gene *LsMAX1* was also induced, rising from baseline at day 0 to approximately five-fold by day 5 and remaining elevated through the remainder of the storage period. *LsMAX2* is the lettuce ortholog of *ORE9*, the founding Arabidopsis stay-green locus first identified in classic forward genetic screens [5, 6], and later shown to encode *MAX2* [7]. Consistent with this link, *LsMAX2* was strongly induced during lettuce storage, rising to approximately seven-fold by day 5 and remaining elevated thereafter (Figure 1A). To test whether SL signaling is sufficient to promote senescence, we treated excised lettuce leaves with the synthetic SL analog racemic GR24 or solvent control and incubated them in darkness. GR24 treatment reduced chlorophyll content by approximately 70% relative to mock treated controls (Figure 1B, Supplementary Figure 1A), demonstrating that exogenous SL is sufficient to strongly accelerate senescence in lettuce. Because racemic GR24 can activate both D14 and KAI2 dependent pathways [42], we repeated this treatment using the single active enantiomer GR24^5DS^, which specifically activates the D14 dependent pathway. This treatment produced the same senescence-accelerating effect as the racemic mixture, supporting a promotive role for canonical D14-dependent SL signaling in lettuce leaf senescence, independent of the KAI2 pathway validation of *LsD14* or *LsMAX2*.Together, these results show that SL pathway components are transcriptionally induced during lettuce postharvest senescence, and that SL signaling through the D14-dependent pathway is sufficient to accelerate it.

**Figure 1.**
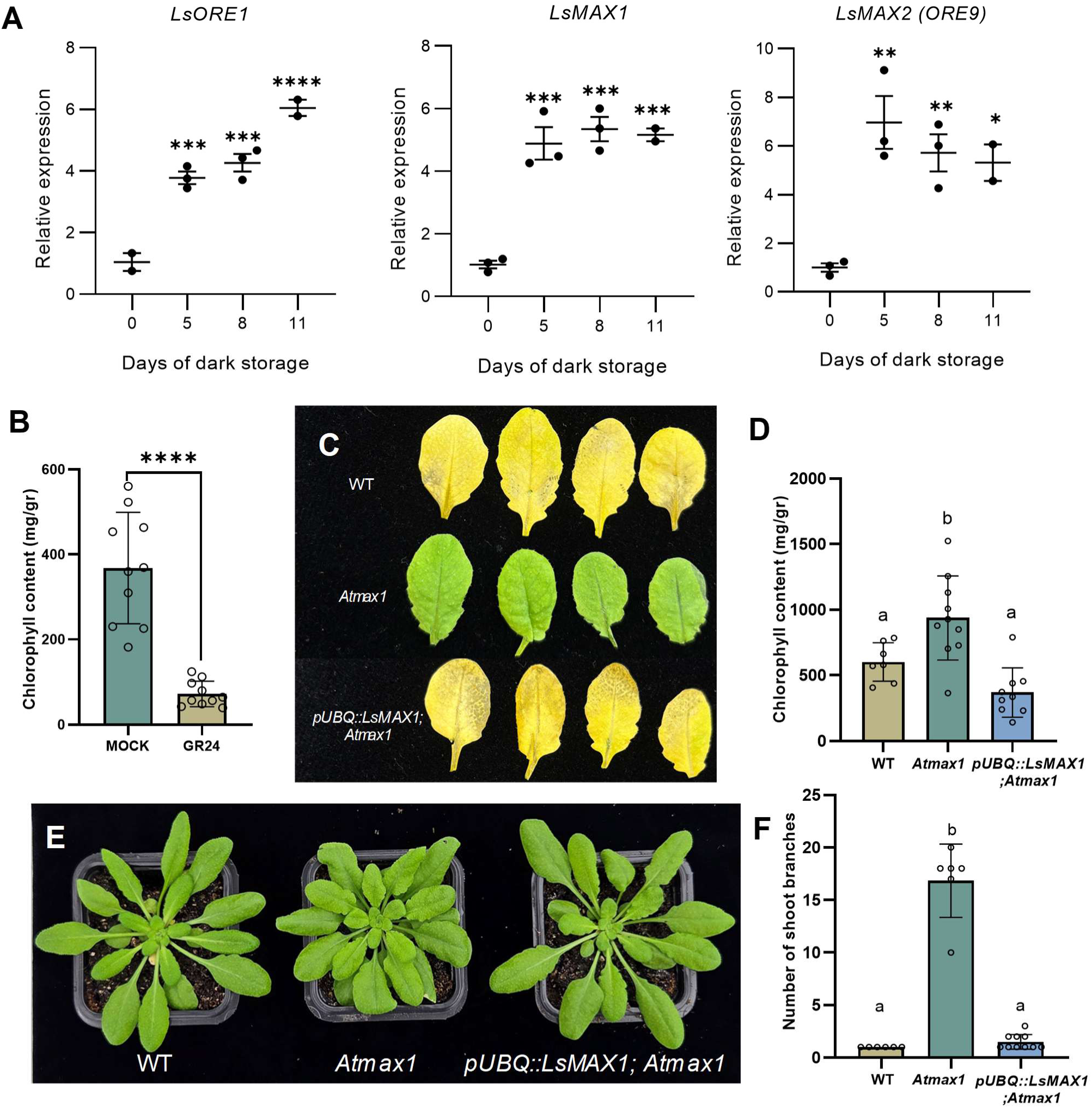
Functional conservation of MAX1 across lettuce and *Arabidopsis thaliana*. A. Relative expression of LsORE1, LsMAX1 and LsMAX2 during lettuce storage, measured by RT-qPCR at days 0, 5, 8, and 11. Data are hown as mean (± s.e.m.), n=2-3 per timepoint. One-way ANOVA with post hoc test; asterisks indicate significant differences (*p<0.05, **p<0.01, ***p<0.001, ****p<0.0001). B. Mean (± s.e.m.) chlorophyll content of lettuce leaves ollowing dark incubation, n=10, t-test, *p*<0.0001. C. Detached leaves from 5-week-old *Arabidopsis thaliana* plants WT, *Atmax1*, and *pUBQ::LsMAX1; Atmax1*) following dark incubation. D. Mean (± s.e.m.) chlorophyll content of leaves of WT (n=7), *Atmax1* (n=10) and *pUBQ::LsMAX1;Atmax1* (n=9), following dark incubation. One-way ANOVA and post hoc tukey test, *p*<0.05. E. Representative images of 5-week-old plants of WT, *Atmax1*, and *pUBQ::LsMAX1; Atmax1*. F. Mean (± s.e.m.) number of axillary rosette branches in WT (n = 6), *Atmax1* (n = 6), and *pUBQ::LsMAX1; Atmax1* (n = 10). One-way ANOVA with Tukey’s post hoc test, *p*<0.0001.

### LsMAX1 encodes a functionally conserved MAX1 enzyme

MAX1 catalyzes a key oxidative step in SL biosynthesis and is present as a single-copy gene in lettuce, making *LsMAX1* the primary target for functional analysis. To confirm that *LsMAX1* encodes a functionally conserved enzyme, we introduced *pUBQ::LsMAX1* into the Arabidopsis *max1-1* mutant, which exhibits delayed dark-induced senescence and enhanced shoot branching, and analyzed T3 homozygous plants from three independent transgenic lines. Expression of *LsMAX1* fully restored both phenotypes: complemented lines senesced at rates indistinguishable from wild type (WT), and total chlorophyll quantification confirmed WT-like degradation kinetics (Figure 1C-D). Rosette morphology and the significantly elevated shoot branch number characteristic of *max1-1* were also fully rescued to WT levels (Figure 1E-F), demonstrating that the lettuce *MAX1* ortholog is sufficient to reinstate canonical SL-dependent developmental processes in Arabidopsis.

### CRISPR/Cas9-mediated disruption of LsMAX1 delays dark-induced senescence in lettuce

To investigate *LsMAX1* function directly in lettuce, we generated loss-of-function alleles using CRISPR/Cas9 in *Lactuca sativa* cv. Salinas. Guide RNAs targeting the first exon produced indel mutations in independent T1 lines, yielding three distinct alleles (Supplementary Figure 2A). The *Lsmax1-1* and *Lsmax1-3* alleles carried small insertions causing frameshifts and premature stop codons, the 10-bp deletion in *Lsmax1-2* similarly caused a frameshift and early termination (Supplementary Figure 2A, red bases). All three alleles are therefore predicted to represent strong loss-of-function variants and were advanced to T3 for phenotypic analysis.

At the young rosette stage, *Lsmax1* seedlings were morphologically indistinguishable from WT, with no differences in rosette architecture, leaf form, or leaf area (Figure 2A-B). Chlorophyll fluorescence (Fv/Fm) measurements in fully expanded leaves also detected no significant differences between WT and *Lsmax1* plants (Figure 2C), indicating that *LsMAX1* is not required for early vegetative growth or baseline photosynthetic efficiency under standard conditions.

**Figure 2.**
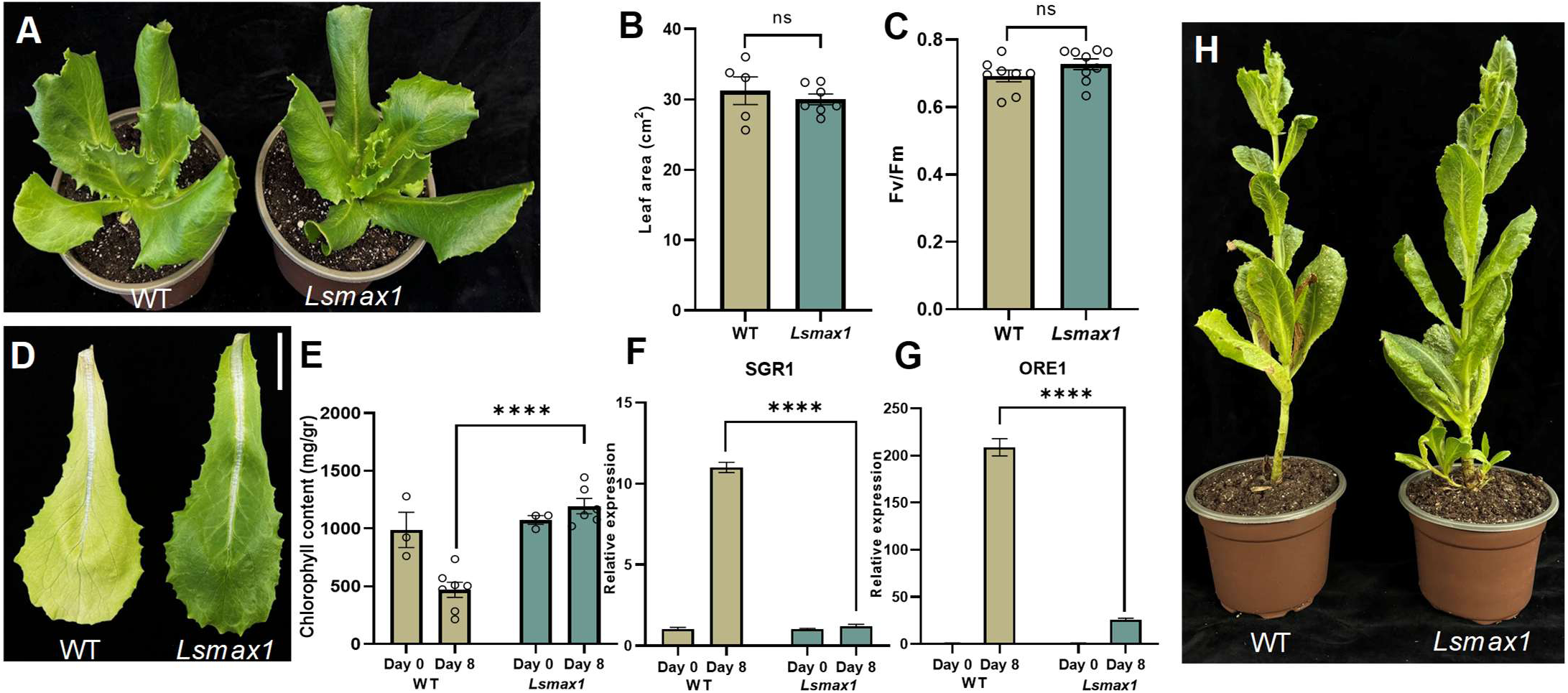
Loss of *LsMAX1* delays senescence in lettuce. **A**. Top view of 6-week-old WT and *Lsmax1* plants showing similar growth and loaf size. **B**. Leaf area of the fourth leaf in WT (n = 5) and *Lsmax1* (n = 7) plants, quantified using ImageJ. *t*-test, not significant (ns) **C**. Fv/Fm (maximum quantum efficiency of PSII; variable fluorescence/maximum fluorescence) of WT and *Lsmax1* plants grown under identical greenhouse conditions, measured using a JUNIOR-PAM fluorometer. *t*-test, not significant (ns). **D**. WT and *max1* Lettuce leaf after 8 days of dark storage. Scale bar = 2cm **E.** Mean (± s.e.m.) chlorophyll content of WT and *Lsmax1* at day 0 (WT, n=3; *Ls*max1, n=3; t-test, ns) and day 8 (WT, n=7; *Lsmax1*, n=6; t-test, *p* < 0.0001). **F-G.** Relative expression levels of senescence gene *LsSGR1* and *LsORE1* in WT and Ls*max1* leaves following dark incubation. Gene expression was quantified using the 2^-^ΔΔCt^ method, and normalization was performed using *ACT2* as a reference gene. Student’s *t*-test, *p*<0.0001, n=3**. H. 6** months old WT and *Lsmax1* flowering plants, illustrating the absence of branching in WT and the presence of axillary branching in *Lsmax1*.

To determine whether SL biosynthesis contributes to senescence, we evaluated *Lsmax1* leaves using two complementary approaches: detached leaves incubated in darkness, and intact leaves darkened on the plant by foil covering. Importantly, chlorophyll content was not significantly different between WT and *Lsmax1* leaves at day 0 prior to dark treatment (Figure 2E), ruling out any developmental difference in chlorophyll accumulation as a confounding factor. After eight days of dark incubation, however, *Lsmax1* leaves remained visibly greener and retained significantly higher chlorophyll content than WT controls in both experimental systems (Figure 2D; Supplementary Figure 2B), demonstrating a pronounced delay in senescence. At the molecular level, the senescence-associated gene *LsSGR1* was induced approximately ten-fold in WT leaves over eight days of dark storage, while this induction was almost completely abolished in *Lsmax1* leaves (Figure 2G). Consistent with its role as a senescence marker established above (Figure 1A), *LsORE1* showed an even more striking genotype-dependent response: expression increased approximately 150- to 200-fold in WT leaves but remained at near-basal levels in *Lsmax1* leaves (Figure 2G). Together, these data demonstrate that *LsMAX1*-dependent SL biosynthesis is required for timely dark-induced senescence in lettuce. The near-complete suppression of *LsSGR1* and *LsORE1* induction in *Lsmax1* leaves indicates that *LsMAX1-*dependent SL biosynthesis is required for their normal induction during dark-induced senescence.

### LsMAX1 loss does not alter vegetative rosette architecture but increases post-bolting branching

We report for the first time that SL-dependent suppression of axillary bud outgrowth is conserved in lettuce, and that this effect is strictly confined to the post-bolting stage. Loss of apical control of axillary meristems was not apparent during early rosette growth (Figure 2A) but became evident only after the transition to flowering, where *max1* plants produced multiple axillary shoots emerging from the main stem, in contrast to the single unbranched stem characteristic of WT plants (Figure 2H). The confinement of architectural changes to the post-bolting stage means that the vegetative rosette structure that determines marketable yield in leafy greens remains fully intact in *Lsmax1* mutants.

### Genotype-specific induction is enriched for known SAGs in the shared response, but not in *Lsmax1*

To determine whether the transcriptional response shared between WT and *Lsmax1* reflects genuine senescence biology, and whether the *Lsmax1*-specific response is drawn from the same canonical gene set, we performed RNA-seq on WT and *Lsmax1* leaves before and after 8 days of dark storage (three biological replicates per genotype and timepoint). *LsMAX1* itself and other core strigolactone biosynthetic genes were not detected among expressed transcripts in this dataset, consistent with their characteristically low abundance. Their *MAX1*-dependent regulation is instead established directly by the qPCR and CRISPR phenotyping data above.

We intersected genes induced during storage in each genotype with a curated reference set of 2,012 Arabidopsis SAG orthologs from the Leaf Senescence Database [43] (Figure 3A). This identified 273 WT-specific and 120 *Lsmax1*-specific SAGs, together with 466 SAGs induced in both genotypes. Genes induced in both genotypes were significantly enriched for canonical SAGs relative to the genome-wide background (1.56-fold enrichment over the 21.3% background rate, hypergeometric test, p = 8.5 × 10⁻³⁰), indicating that the shared transcriptional core reflects genuine senescence biology rather than incidental overlap between genotypes with markedly different visible phenotypes. Genes induced specifically in WT were similarly enriched (1.40-fold, p = 1.1 × 10⁻¹⁰). Genes induced specifically in *Lsmax1*, in contrast, showed no such enrichment (1.07-fold, p = 0.199), indicating that this genotype-specific response is not drawn from the canonical, literature-recognized senescence gene set.

**Figure 3.**
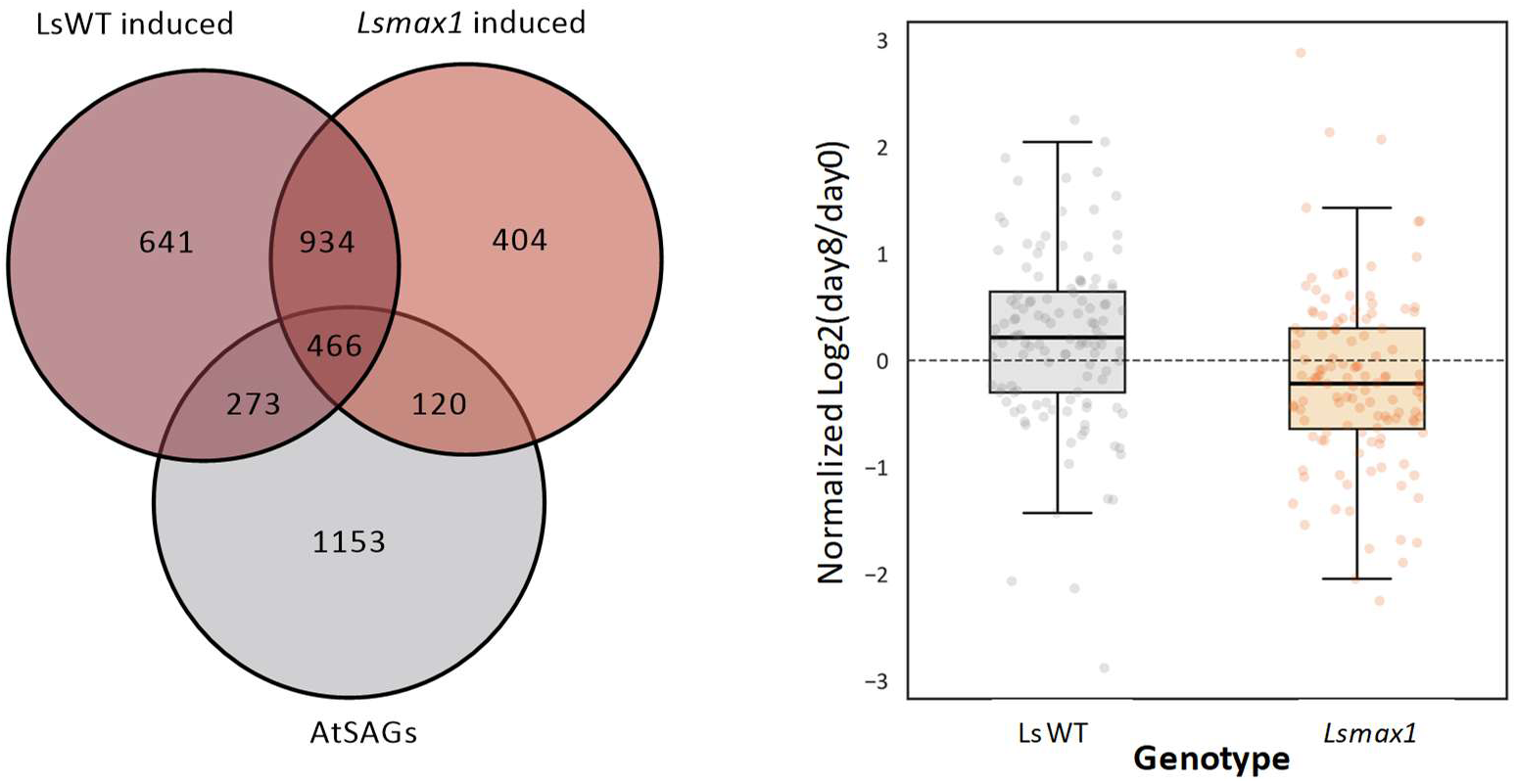
Genotype-specific induction is enriched for known SAGs in the shared response, but not in *Lsmax1*. A. Venn diagram showing the overlap between genes induced in WT and *Lsmax1* during dark storage and a curated reference set of 2,012 Arabidopsis senescence-associated gene (SAG) orthologs from the Leaf Senescence Database (LSD v5.0, Zhao et al., 2024; “AtSAGs”). Genes induced in both genotypes are significantly enriched for canonical SAGs relative to the genome-wide background (1.56-fold, hypergeometric test, p = 8.5 × 10⁻³⁰), as are genes induced specifically in WT (1.40-fold, p = 1.1 × 10⁻¹⁰), whereas genes induced specifically in *Lsmax1* show no such enrichment (1.07-fold, p = 0.199). B. Normalized log2 fold change (day 8 vs. day 0) in WT and *Lsmax1* across an independent, literature-curated set of 120 Arabidopsis genes annotated as upregulated during senescence (181 lettuce orthologs; paralogs averaged per Arabidopsis gene to preserve independence of observations). WT and *Lsmax1* age similarly at the transcriptional level, with only minor differences detected between genotypes

To test this pattern using a fully independent, literature-curated gene set rather than genes selected by significance in our own data, we compiled 365 Arabidopsis genes explicitly annotated as upregulated during senescence in the Leaf Senescence Database. Of these, 120 mapped to lettuce orthologs (181 lettuce genes in total; for the 37 Arabidopsis genes with multiple lettuce paralogs, fold changes were averaged across paralogs to preserve independence of observations). Because the distribution of WT-*Lsmax1* differences departed significantly from normality (Shapiro-Wilk test, p = 0.0131), genotypes were compared using a Wilcoxon signed-rank test. WT and *Lsmax1* log2 fold-change values (day 8 vs. day 0) differed significantly across this independent gene set (Wilcoxon signed-rank test, p = 0.0031) (Figure 3B), with WT showing a modestly positive median shift and *Lsmax1* centered close to zero. Because relatively few genes in the database were explicitly annotated as downregulated during senescence, this analysis was restricted to the independently curated set of upregulated SAGs. This independent analysis reinforces the conclusion that WT and *Lsmax1* retain broadly similar senescence-associated transcriptional responses during dark storage, with only modest differences in canonical SAG activation despite their markedly different outward phenotypes.

### WT and *Lsmax1* share a common transcriptional core but diverge in senescence activation

At the genome-wide level, WT and *Lsmax1* shared a substantial fraction of their transcriptional response during storage. 1,400 of 2,838 upregulated genes (49.3%) and 2,125 of 3,539 downregulated genes (60.0%) were concordantly regulated in both genotypes (Figure 4A). This asymmetry, in which a larger fraction of repressed genes is shared than of induced genes, indicates that WT and *Lsmax1* diverge more in which genes they activate than in which genes they shut down, even though a large core of both responses is shared. GO enrichment of the concordantly regulated genes reinforced this pattern (Figure 4B). The shared repressive and shutdown component is dominated by strong, canonical senescence categories (photosynthesis, translation), whereas the shared inductive component shows comparatively weak and functionally uncharacterized enrichment. Together, these results indicate that WT and *Lsmax1* leaves both undergo senescence, but through partially distinct transcriptional routes, with *Lsmax1* engaging a smaller and less coherent activation program than WT.

**Figure 4.**
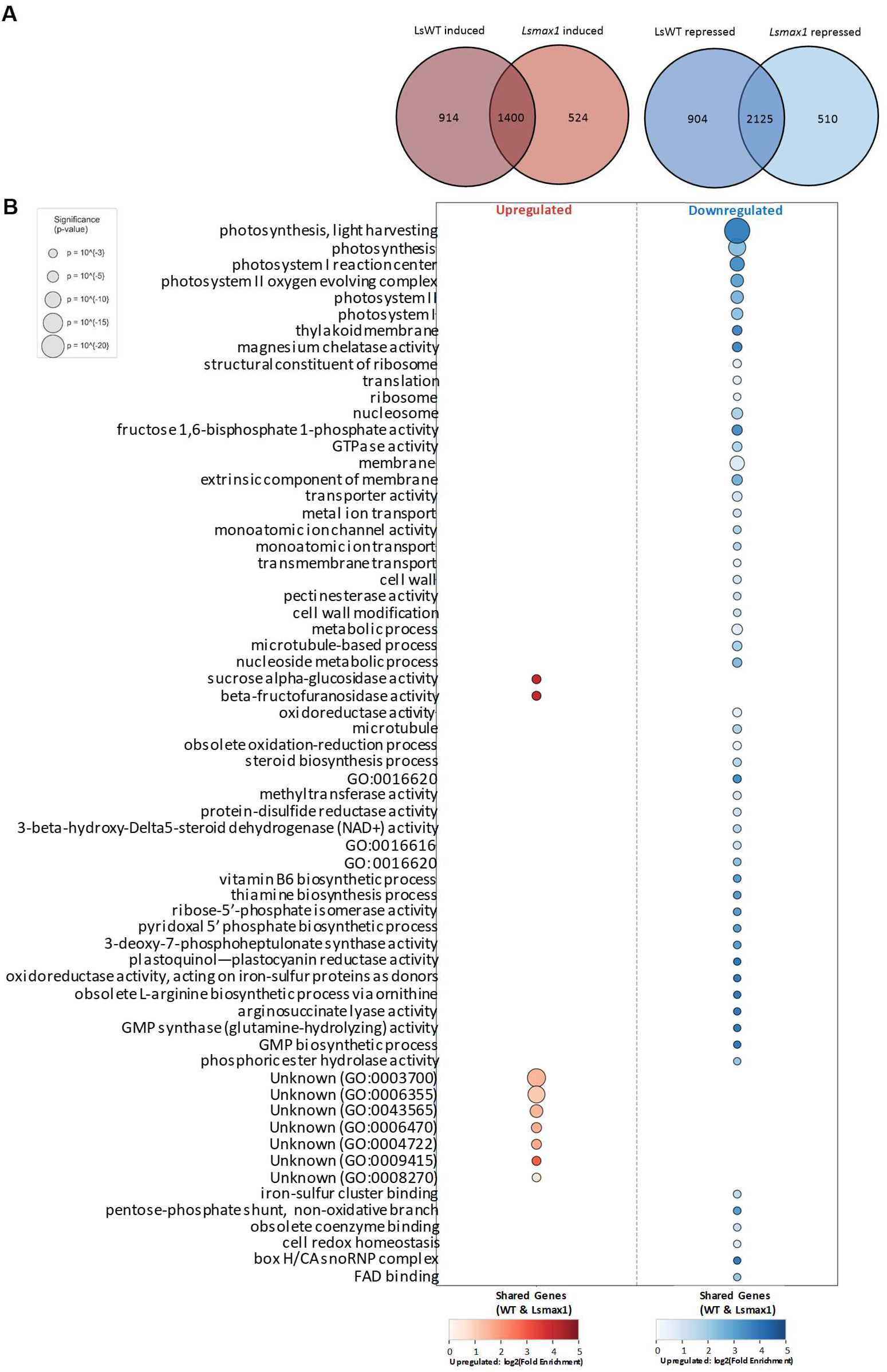
WT and *Lsmax1* share a common transcriptional core but diverge in senescence activation. A. Venn diagrams of genes upregulated (left) and downregulated (right) in WT and *Lsmax1* during dark storage (padj < 0.05). The two genotypes share the majority of their transcriptional response despite *Lsmax1*’s visibly reduced senescence progression, with a larger fraction of repressed genes shared than induced genes. B. GO term enrichment of genes concordantly regulated in both genotypes (shared upregulated and downregulated gene sets from A). The shared repressive and shutdown component is dominated by strong, canonical senescence categories (photosynthesis, translation), whereas the shared inductive component shows comparatively weak and functionally uncharacterized enrichment. Dot size indicates significance (p-value), dot color indicates log2 fold enrichment.

### The *Lsmax1*-specific response defines a distinct stress-associated transcriptional module

Having established that *Lsmax1*-specific induced genes are not enriched for canonical SAGs (Figure 3A), we next asked whether the WT-specific and *Lsmax1*-specific responses, defined by the genome-wide Venn overlaps in Figure 4A, differ functionally. GO term enrichment was performed on the complete genome-wide genotype-specific gene sets (932 WT-specific and 545 *Lsmax1*-specific upregulated genes, and 935 WT-specific and 523 *Lsmax1*-specific downregulated genes) (Figure 5A). Among upregulated genes, WT-specific genes were enriched for chromatin- and transcription-associated categories (nucleosome, RNA polymerase II regulation, mediator complex) together with autophagy, consistent with a canonical, transcriptionally driven senescence program. *Lsmax1*-specific upregulated genes, in contrast, were enriched for stress response, ubiquitin-proteasome, and calcium/calmodulin signaling categories, representing a functionally distinct transcriptional module rather than a weaker copy of the WT program.

**Figure 5.**
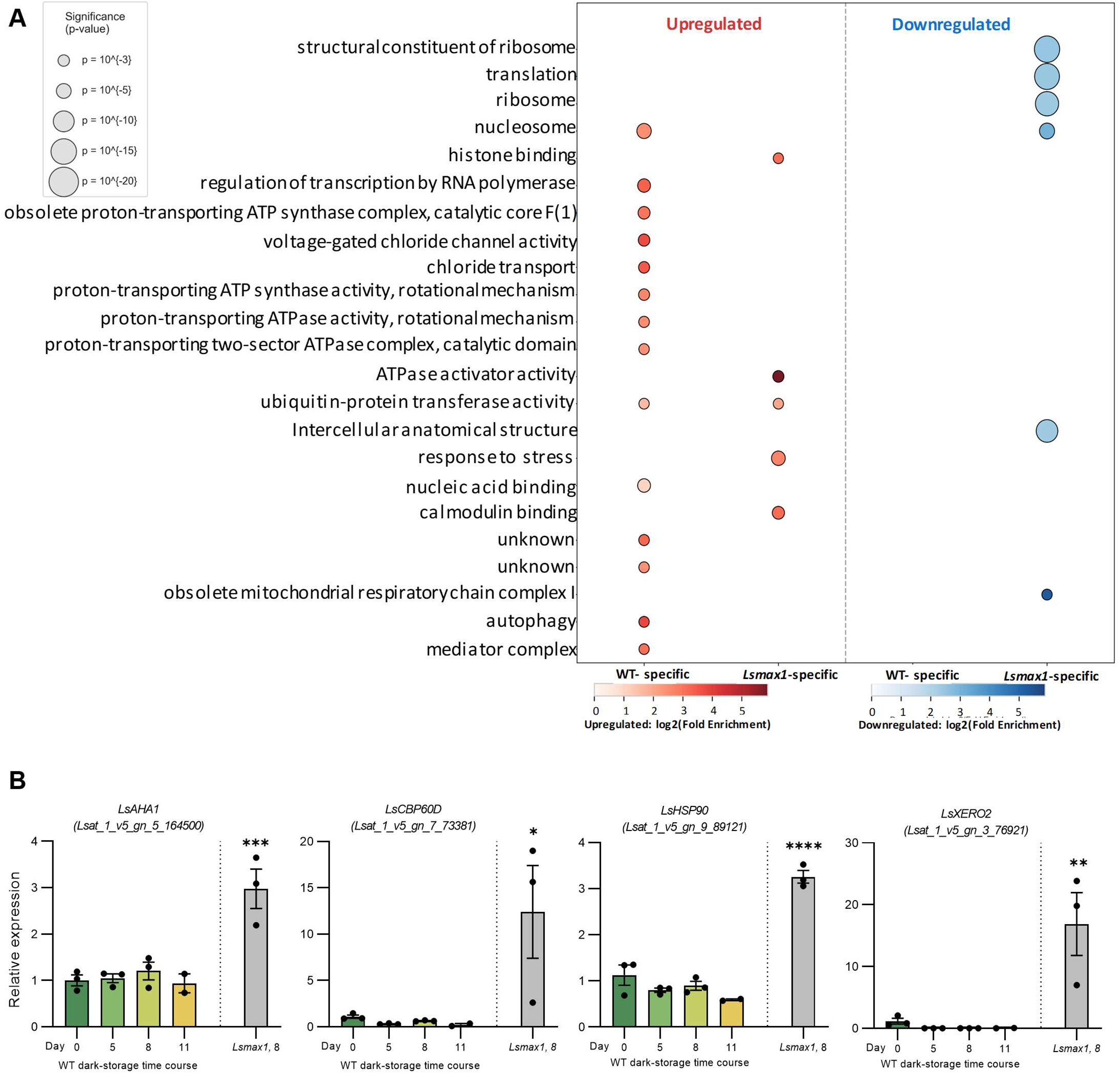
WT- and *Lsmax1*-specific genes engage functionally distinct transcriptional programs. A. GO term enrichment of genes upregulated or downregulated specifically in WT or specifically in *Lsmax1* during dark storage (genome-wide gene sets, not restricted to the SAG reference list). WT-specific genes were enriched for chromatin- and transcription-associated categories nucleosome, RNA polymerase II regulation, mediator complex) together with autophagy. No significant GO enrichment was detected among genes downregulated specifically in WT. *Lsmax1*-specific upregulated genes were enriched for stress response, ubiquitin-proteasome, and calcium/calmodulin signaling categories, while *Lsmax1*-specific downregulated genes were enriched for ribosome- and translation-associated categories. Dot size indicates significance (p-value), dot color indicates log2 fold enrichment.B. RT-qPCR validation of four genes from the *Lsmax1*-specific stress-response module (*LsAHA1*, *LsCBP60D*, *LsHSP90*, *LsXERO2*) across a WT dark-storage time course (days 0, 5, 8, 11) and in *Lsmax1* at day 8 (same RNA samples analyzed by RNA-seq). All four genes were induced in *Lsmax1* while remaining low throughout the WT time course. Data are shown as mean (± s.e.m.); one-way ANOVA with post hoc test, asterisks indicate significant differences relative to WT day 0 (*p<0.05, **p<0.01, ***p<0.001, p<0.0001).

Among downregulated genes, *Lsmax1*-specific genes were enriched for ribosome- and translation-associated categories. No significant GO enrichment was detected among genes downregulated specifically in WT, indicating that the genes uniquely repressed in WT lack a coherent functional signature, in contrast to the *Lsmax1*-specific downregulated set. This *Lsmax1*-specific induction represents a transcriptional program that remains largely unengaged in WT during storage, indicating that loss of *LsMAX1* is associated with activation of an alternative, stress-associated transcriptional response rather than simply reduced activation of the canonical senescence program. To validate genes identified from the *Lsmax1*-specific GO-enriched stress-response category, we measured expression of *LsXERO2*, *LsHSP90*, *LsCBP60D*, and *LsAHA1* by RT-qPCR. In *Lsmax1*, all four genes were induced at day 8 of dark storage, using the same RNA samples analyzed by RNA-seq, consistent with the RNA-seq finding that this gene set is engaged specifically in the mutant. Notably, *LsHSP90* and *LsAHA1*, which encodes the HSP90 co-chaperone Activator of Hsp90 ATPase 1, were induced together. In an independent WT storage time course, expression of these genes remained low throughout, supporting genotype-specific rather than storage-stage-dependent induction (Figure 5B). Together, these results indicate that the *Lsmax1*-specific response is not simply a diminished version of the WT program, but a distinct stress-associated transcriptional module, supported by independent validation of representative genes.

## Discussion

Strigolactones (SLs) are established positive regulators of leaf senescence in Arabidopsis, and several other species [1, 2, 4], yet their endogenous contribution to senescence in leafy vegetable crops has remained unclear. Here, genetic disruption of *LsMAX1* establishes a requirement for endogenous SL biosynthesis in the normal progression of dark-induced senescence in lettuce. Together with the senescence-promoting effect of GR24, including the D14-dependent enantiomer GR24^5DS^ in both lettuce and Arabidopsis (Supplementary Figure 1B, C), and the functional complementation of Arabidopsis *max1-1* by *LsMAX1*, these findings extend the SL-senescence relationship from model systems to a commercially important leafy crop. More importantly, the transcriptional response of *Lsmax1* indicates that its stay-green phenotype cannot be explained simply as a uniform delay of the WT senescence program. Instead, MAX1-dependent SL biosynthesis appears to influence how the senescence transcriptional program is configured under prolonged darkness.

Despite the pronounced difference in visible yellowing, a substantial transcriptional response to dark storage was retained in *Lsmax1*, including a shared core enriched in recognized senescence-associated genes (SAGs). Thus, loss of *LsMAX1* does not abolish the broader molecular response associated with senescence. Instead, the major divergence between genotypes occurs in the genes activated during this response. This asymmetry between transcriptional activation and repression provides a framework for understanding the role of MAX1. Cellular shutdown processes, including repression of photosynthesis and translation-related functions, were prominent within the response shared by WT and *Lsmax1*. By contrast, WT-specific induction was associated with transcriptional and chromatin regulatory functions and autophagy, whereas the *Lsmax1*-specific response was associated with stress signaling, protein homeostasis, ubiquitin-dependent processes, and calcium-related signaling. Moreover, *Lsmax1*-specific induced genes were not enriched for established SAGs. The mutant therefore appears to retain much of the shutdown component of the dark-storage response while replacing part of the WT activation program with an alternative stress-associated transcriptional state.

The strong *LsMAX1* dependence of *LsORE1* and *LsSGR1* induction is consistent with this interpretation. ORE1 is a central transcriptional regulator of senescence in Arabidopsis, while SGR proteins connect senescence signaling with chlorophyll degradation. Their strongly reduced induction in *Lsmax1* links the altered transcriptional program with the persistent green phenotype. However, the genome-wide response indicates that *LsMAX1* does not control only chlorophyll degradation or a small set of terminal senescence genes. Rather, its loss alters a broader activation component while allowing substantial transcriptional remodeling to continue. In this sense, the *Lsmax1* phenotype is better described as a redirection of the senescence program than as its arrest. The independently validated *Lsmax1*-specific stress module further supports this model. Particularly notable is the coordinated induction of *LsHSP90* and *LsAHA1*. HSP90 functions in protein quality control and plant stress responses [44], while AHA1 is an HSP90 co-chaperone that stimulates its ATPase activity [45]. Their coordinated induction is therefore consistent with increased engagement of HSP90-associated chaperone machinery in *Lsmax1*. Whether this response contributes functionally to the stay-green phenotype or instead reflects the altered physiological state of the mutant remains to be determined.

Current models of SL-dependent senescence have largely emphasized interactions between SLs and other hormone pathways. In Arabidopsis, SL signaling interacts with ethylene during dark-induced senescence [1], and more recent work places SL signaling upstream of a sequential SL-to-salicylic-acid pathway that promotes senescence [38]. Our findings suggest an additional level of regulation. Rather than acting only as a quantitative accelerator of a fixed hormone-driven pathway, MAX1-dependent SL biosynthesis may influence which transcriptional modules are recruited during senescence. Loss of this input leaves much of the common response intact but changes the balance between canonical senescence-associated activation and alternative stress responses.

Such a role is particularly plausible under prolonged darkness. SL biosynthesis is closely linked to nutrient and carbon status: phosphate limitation promotes SL production, while extended darkness and sugar depletion induce SL biosynthetic genes, and exogenous sugar can suppress SL-promoted senescence [4, 32–34, 36, 37]. Postharvest lettuce leaves experience sustained carbon limitation because separation from the plant eliminates continued carbon import while darkness prevents photosynthetic carbon assimilation. The induction of *LsMAX1* during storage is therefore consistent with a model in which endogenous SL biosynthesis contributes to converting sustained carbon limitation into a senescence-associated transcriptional program. Direct measurements of SL metabolites and carbon status will be required to test this model.

The distinction between redirection and simple delay may also help reconcile apparently inconsistent effects of SL deficiency on senescence among species. Clear stay-green or delayed-senescence phenotypes have been reported in Arabidopsis, rice, bamboo, and petunia [1, 2, 4, 8], whereas similarly obvious phenotypes have not been described in SL mutants of pea or tomato [9–13]. If SL signaling influences the recruitment of particular senescence-associated transcriptional modules rather than a single universally conserved rate-limiting step, its phenotypic effects could depend on the alternative programs available in different species or environmental contexts.

The developmental restriction of the branching phenotype in lettuce further distinguishes the consequences of manipulating SL biosynthesis in this crop. Suppression of axillary bud outgrowth is one of the best-characterized functions of SLs, and disruption of SL biosynthesis or signaling produces branching or tillering phenotypes in species including Arabidopsis, rice, pea, and tomato [9, 11, 33]. In lettuce, however, enhanced branching became apparent only after bolting. Because commercial lettuce is harvested during the vegetative rosette stage, modification of SL biosynthesis may therefore influence leaf longevity without substantially altering the architecture of the harvested plant.

An important limitation is that WT and *Lsmax1* were compared at the same chronological time but at visibly different stages of yellowing. Some genotype-specific expression differences may therefore reflect differences in physiological progression rather than direct transcriptional targets of *MAX1*-dependent signaling. Nevertheless, the extensive shared response, absence of SAG enrichment among *Lsmax1*-specific induced genes, and independent validation of mutant-specific stress-associated genes argue against a simple global developmental offset. Time-resolved transcriptomics, ideally combined with SL treatment or physiological stage matching, will be necessary to distinguish primary *MAX1*-dependent events from downstream consequences of altered senescence progression. Identification of endogenous SL species and genetic analysis of *LsD14* and *LsMAX2* will also be important next steps.

Overall, our findings identify *MAX1*-dependent SL biosynthesis as an important determinant of leaf senescence in lettuce and suggest a role beyond simply controlling its rate. In the absence of *LsMAX1*, leaves retain a substantial senescence-associated transcriptional core but fail to fully engage the WT activation program and instead recruit a distinct stress-associated response. *MAX1*-dependent SL biosynthesis may therefore help determine the transcriptional trajectory adopted during prolonged darkness. Because this stay-green phenotype occurs without detectable disruption of vegetative rosette architecture, this pathway may also provide a target for extending postharvest longevity in lettuce and related leafy crops.

## Materials and methods

### Plant growth and leaf storage conditions

Seeds of *Lactuca sativa* L. cv. Salinas were obtained from the laboratory of Richard W. Michelmore (University of California, Davis, CA, USA). The *Lsmax1* mutant was generated in the *L. sativa* cv. Salinas background as described below. Seeds of *Arabidopsis thaliana* (Col-0 background) were obtained from the Arabidopsis Biological Resource Center (ABRC), including CS9564 (*max1-1*). No plant material used in this study was collected from the wild. All experimental research on plants was conducted in accordance with relevant institutional, national, and international guidelines.

Lettuce (*L. sativa* cv. Salinas, WT) and *Lsmax1* mutant plants were transplanted into individual square pots (9 x 9 x 5.5 cm) containing a peat-perlite potting mix and grown in a greenhouse at 23C under natural light supplemented with artificial lighting from 08:00 to 22:00. After one month, plants were transferred to larger pots (10.5 x 11 x 11.5 cm) for continued growth. For detached-leaf dark-induced senescence assays, the fourth fully expanded leaf from the base of WT and *Lsmax1* 5-week-old plants was excised and stored in high-density polyethylene bags to maintain 95-98% relative humidity. Leaves were kept at room temperature in complete darkness as described by [46]. At least four biological replicates per genotype were used. Leaves were photographed and chlorophyll content was quantified as described below. For attached-leaf dark-induced senescence assays, the fourth fully expanded leaf was individually wrapped in aluminum foil to exclude light while remaining attached to the plant. Plants were maintained under standard greenhouse conditions (23C, supplemented lighting from 08:00 to 22:00) for 14 days. At least four biological replicates per genotype were used. Leaves were photographed and chlorophyll content was quantified as described below. For Arabidopsis assays, fully expanded rosette leaves were detached from 5-week-old WT, *Atmax1*, and *pUBQ::LsMAX1*;*Atmax1* plants. Detached leaves were placed on moistened filter paper in sealed Petri dishes and incubated in complete darkness at 25C for 7 days. At least four biological replicates per genotype were used. Leaves were photographed and chlorophyll content was quantified as described below.

### GR24 treatment

Detached lettuce leaves were treated with 10 uM racemic GR24 (Chiralix, Product No. CX23880) or the single active enantiomer GR24^5DS^ (BOC Sciences, Product No. 151716-22-2) and acetone was used as the solvent control. The leaves were stored in the dark at 25C. Leaves were weighed before chlorophyll extraction, and chlorophyll was extracted in 1 mL DMF. Chlorophyll was measured spectrophotometrically at 647 and 664 nm. Chlorophyll concentration per gram of leaf tissue was calculated using the formula, chlorophyll concentration (mg per g) = (20.27 x A647) + (7.04 x A664) [47].

### Phylogenetic analysis

Phylogenetic analysis was performed using the Custom Phylogenetic Tree tool in the PLAZA Dicots platform [48]. Protein homologs were identified using BLASTP against the PLAZA Dicots database, using the *Arabidopsis thaliana* sequence as the query, and including homologs from Lactuca sativa, *Arabidopsis thaliana*, Solanum lycopersicum, Quercus lobata, Olea europaea, and Citrus clementina. Tree visualization and final figure design were performed using the iTOL website [49].

### Complementation assay in Arabidopsis

Golden Gate Modular Cloning Toolbox for Plants (MoClo) [50] was used to overexpress MAX1 in Arabidopsis. The coding sequence of *LsMAX1* (Lsat_1_v5_gn_6_14580; NCBI RefSeq accession XM_023881386.3; protein accession XP_023737154.1), corresponding to *Lactuca sativa* cytochrome P450 711A1 (CYP711A1), was custom-synthesized with suitable restriction enzyme recognition and cleavage sites and appropriate overhangs for MoClo assembly. *LsMAX1* was cloned into the pICH41308 plasmid (Level 0) using the BbsI restriction enzyme. The Level 0 module was cloned into the pICH47742 plasmid (Level 1) with the UBQ10 promoter and NOS terminator using the BsaI restriction enzyme. The Level 1 module was cloned into the pAGM4723 plasmid (Level 2) for transformation of *Arabidopsis thaliana*. This binary vector served as the backbone for constructing the multigene system, and the kanamycin-resistance gene (NPTII) was cloned into it to enable selection of transgenic plants. The final construct was introduced into Agrobacterium electrocompetent cells (GV3101) and subsequently transformed into the *max1-1* insertion line, ABRC stock CS9564. T1 seeds were harvested and sown on 0.5 MS agar medium containing 50 uM kanamycin. The resistant seedlings were transferred to soil for growth until T2 seeds were harvested. We obtained three independent T3 homozygous complementation lines.

### RNA extraction and qPCR

The fourth leaf of WT and *Lsmax1* plants was stored in a zip-bag under dark conditions at 25C for 8 days. RNA was extracted using TRIzol (TRIzol Reagent, Thermo Scientific, catalog number 15596026) using 50 to 100 mg tissue per 1 mL TRIzol. 1 ug RNA was then treated with a DNase kit (DNase I, RNase-free, Thermo Scientific, catalog number EN0521). RNA was reverse-transcribed into cDNA using the LunaScript RT SuperMix Kit (New England Biolabs, catalog number NEB-M3010L). To assess the progress of dark-induced leaf senescence during storage, the dynamic expression of ORE1 and SGR1, a senescence-associated gene, was monitored using quantitative real-time PCR (qPCR) with a QuantStudio 1 system (Applied Biosystems). qPCR reactions were prepared using a SYBR mix (Bio-Rad, iTaq Universal SYBR Green Supermix, catalog number 1725121). Each experiment was performed using three independent biological replicates, each analyzed in three technical replicates. Relative gene expression was calculated using the 2^-ddCt method, with normalization to the lettuce ACT2 reference gene.

### RNA-seq and differential expression analysis

WT and *Lsmax1* leaves were sampled at day 0 and day 8 of dark storage, three biological replicates per genotype per time point. Sequencing libraries were prepared using MARS-seq. Single-end libraries were sequenced on an Illumina NovaSeq platform. The output was ∼15 million reads per sample. Raw reads were trimmed with Cutadapt v5.2 [51] and aligned to the Lactuca sativa reference genome (Lsativa_467 v8, annotation v5 [52] using STAR (v2.7.11b [53]). PCR duplicates were identified and removed using Unique Molecular Identifier (UMI) barcoding (UMI-tools v1.1.4 [54]) prior to gene-level quantification with featureCounts (v2.1.1 [55]). Differential expression was assessed in R using DESeq2 (v1.52.0 [56] using a negative binomial model with library-size normalization. Genes with an adjusted *P* value (padj) < 0.05 and an absolute log2 fold change (|log2FC|) ≥ 1 were considered differentially expressed.

### Enrichment of senescence-associated genes among genotype-specific responses

To test whether genes induced specifically in each genotype are enriched for canonical senescence-associated genes (SAGs), we performed a hypergeometric test against a background of 9,427 testable lettuce genes, of which 2,012 are annotated SAG orthologs in the Leaf Senescence Database (LSD v5.0, [43]). Enrichment was calculated separately for genes induced in both genotypes and for genes induced specifically in *Lsmax1*, using the observed and expected number of SAGs in each set.

### Comparison of WT and Lsmax1 expression across an independent curated SAG set

To test the shared senescence response using a fully independent gene set, we identified 365 Arabidopsis genes annotated as upregulated during senescence in the Leaf Senescence Database (LSD v5.0, [43]). Because relatively few genes in the database were explicitly annotated as downregulated during senescence, downregulated SAGs were not included in this analysis. Of the 365 genes, 120 had lettuce orthologs represented in our RNA-seq dataset, corresponding to 181 lettuce genes in total. For the 37 Arabidopsis genes with more than one lettuce paralog, log2 fold change values were averaged across paralogs to yield one independent value per Arabidopsis gene. Fold change values (day 8 vs. day 0) were normalized by centering each gene’s WT and *Lsmax1* values around their shared mean. Normality of the WT-*Lsmax1* differences was assessed using the Shapiro-Wilk test, which rejected the normality assumption (p = 0.0131). Genotypes were therefore compared using a two-sided Wilcoxon signed-rank test.

### Cloning of plasmids for CRISPR/Cas9

To clone two gRNA expressing cassettes and minimize repetition, we modified pMR217 [57] and replaced the AtU6-26 promoter with a synthetic promoter based on the consensus sequence of the three U6 promoter variants present in the Arabidopsis genome [58]. In addition, we modified the scaffold of the sgRNA according to [59]. We used CRISPOR [60] to select two gRNAs targeting *LsMAX1*, ranked by predicted efficiency according to [61]. The resulting sgRNA cassettes were transferred by LR recombination into the pMR924 binary vector, which contains LsUBIp-Cas9 and a green fluorescent protein visible marker, to generate pMR929. This vector was subsequently introduced into A. tumefaciens strain LBA4404 for stable transformation.

### Agrobacterium-mediated transformation and regeneration of lettuce

*L. sativa* cv. Salinas was transformed using a modified cotyledon-based Agrobacterium tumefaciens protocol [62]. Seeds were surface-sterilized in 20% bleach for 20 min, rinsed extensively with sterile water, and germinated on half-strength Hoagland’s medium. Seedlings were grown for 4 days at 24C under a 12 h light and 12 h dark photoperiod. Cotyledon explants were excised and incubated in A. tumefaciens suspension, followed by co-cultivation on Schenk and Hildebrandt (SH) medium supplemented with acetosyringone, 6-benzylaminopurine, and 1-naphthaleneacetic acid for 3 days in the dark. Explants were then transferred to SH induction medium containing 6-BAP, 1-NAA, and antibiotics for selection and bacterial suppression. Regenerated shoots were transferred sequentially to shoot elongation and rooting media based on SH or Murashige and Skoog formulations, with periodic subculturing until robust shoot and root development. Rooted plantlets were acclimated to soil under high humidity and subsequently transferred to greenhouse conditions, where they were grown to maturity and inflorescences were bagged to prevent seed loss and cross pollination. T1 plants were genotyped by PCR amplification and sequencing of the MAX1 locus to confirm mutant alleles. T2 plants were grown to maturity and T3 seeds were harvested from individual T2 lines confirmed to carry mutant alleles. T3 plants were genotyped by PCR amplification and Sanger sequencing of the *LsMAX1* locus to confirm homozygosity at all three alleles prior to phenotypic analysis.

### Chlorophyll fluorescence (Fv/Fm) measurements

Leaves from WT and *Lsmax1* plants were analyzed using a Junior-PAM chlorophyll fluorometer. Before measurement, plants were dark-adapted for 15 minutes. Measurements were taken immediately after dark adaptation on fully expanded leaves at a consistent position on the leaf blade, avoiding major veins. Fv/Fm was calculated from Fo and Fm values obtained using a saturating pulse, and multiple biological replicates were measured for each genotype under the same conditions.

### Leaf area quantification

Leaf area was quantified in the fourth leaf of WT (n = 5) and *Lsmax1* (n = 7) plants at the rosette stage. Leaves were detached, placed on a flat surface alongside a ruler for scale, and scanned. Leaf area was measured from digital images using ImageJ software (National Institutes of Health, USA) by tracing the leaf outline and applying the scale calibration.

### Shoot branch quantification

Shoot branching was quantified in WT and *Lsmax1* plants at approximately 60 days after germination, at which point WT and mutant plants were at the same developmental age. For Arabidopsis, axillary rosette branches longer than 5 mm were counted in 5-week-old plants of WT, *Atmax1*, and *pUBQ::LsMAX1*;*Atmax1*.

## Statistical analysis

All statistical analyses were performed using GraphPad Prism. Differences between two groups were analyzed using Welch’s t-test, while comparisons among multiple groups were conducted using one-way analysis of variance (ANOVA). One-way ANOVA was followed by Tukey’s multiple comparison test. A p-value of p < 0.05 was considered statistically significant.

## Abbreviations

ABRC: Arabidopsis Biological Resource Center
ANOVA: Analysis of variance
CRISPR: Clustered regularly interspaced short palindromic repeats
DMF: N,N-Dimethylformamide
GO: Gene Ontology
LSD: Leaf Senescence Database
MAX1: MORE AXILLARY GROWTH1
MoClo: Modular Cloning
NCBI: National Center for Biotechnology Information
RNA-seq: RNA sequencing
RT-qPCR: Reverse transcription quantitative PCR
SAG: Senescence-associated gene
SL: Strigolactone
SRA: Sequence Read Archive
UMI: Unique molecular identifier
WT: Wild type.

## Declarations

### Ethics approval and consent to participate

Not applicable.

### Consent for publication

Not applicable.

### Availability of data and materials

The RNA-seq datasets generated during the current study are available in the European Nucleotide Archive (ENA) under project accession number PRJEB124228. All other data generated or analyzed during this study are included in this article and its supplementary information files. Seeds and plant materials generated in this study are available from the corresponding author upon reasonable request.

## Author contributions

A.K. and G.A. performed experiments, with A.K. carrying out the majority of the experimental work. N.O. and D.R. performed the bioinformatics analyses, with D.R. additionally providing guidance and supervision to N.O. Mily Ron performed cloning, plant transformation, and cultivation of the first generation of transformed plants, and Megan Reeves performed plant transformation. R.M. provided scientific advice. A.K. and L.T. wrote the manuscript, with critical feedback and manuscript review from D.R. and R.M. All authors reviewed and approved the final manuscript.

## Acknowledgements

The authors thank the members of the Tal, Russ, and Michelmore laboratories for helpful discussions, scientific input, and for creating a supportive and collaborative research environment throughout this work.

## Declaration of generative AI and AI-assisted technologies in the manuscript preparation process

During the preparation of this work the authors used Claud and Chat GPT for structural assistance and improving readability. After using these tools, the authors reviewed and edited the content as needed and take full responsibility for the content of the published article.

## References

1. Ueda H, Kusaba M. Strigolactone Regulates Leaf Senescence in Concert with Ethylene in Arabidopsis. Plant Physiol. 2015;169:138–47.

2. Yamada Y, Furusawa S, Nagasaka S, Shimomura K, Yamaguchi S, Umehara M. Strigolactone signaling regulates rice leaf senescence in response to a phosphate deficiency. Planta. 2014;240:399–408.

3. Yamada Y, Umehara M. Possible Roles of Strigolactones during Leaf Senescence. Plants. 2015;4:664–77.

4. Tian M-Q, Jiang K, Takahashi I, Li G-D. Strigolactone-induced senescence of a bamboo leaf in the dark is alleviated by exogenous sugar. J Pestic Sci. 2018;43:173–9.

5. Oh SA, Park JH, Lee GI, Paek KH, Park SK, Nam HG. Identification of three genetic loci controlling leaf senescence in Arabidopsis thaliana. Plant Journal (United Kingdom). 1997.

6. Woo HR, Chung KM, Park JH, Oh SA, Ahn T, Hong SH, et al. ORE9, an F-box protein that regulates leaf senescence in Arabidopsis. Plant Cell. 2001;13:1779–90.

7. Stirnberg P, van De Sande K, Leyser HMO. MAX1 and MAX2 control shoot lateral branching in Arabidopsis. Development. 2002;129:1131–41.

8. Snowden KC, Simkin AJ, Janssen BJ, Templeton KR, Loucas HM, Simons JL, et al. The Decreased apical dominance1/Petunia hybrida CAROTENOID CLEAVAGE DIOXYGENASE8 gene affects branch production and plays a role in leaf senescence, root growth, and flower development. Plant Cell. 2005;17:746–59.

9. Beveridge CA, Symons GM, Murfet IC, Ross JJ, Rameau C. The rms1 Mutant of Pea Has Elevated Indole-3-Acetic Acid Levels and Reduced Root-Sap Zeatin Riboside Content but Increased Branching Controlled by Graft-Transmissible Signal(s) on JSTOR. Plant Physiol. 1997;115:1251–8.

10. Morris SE, Turnbull CG, Murfet IC, Beveridge CA. Mutational analysis of branching in pea. Evidence that Rms1 and Rms5 regulate the same novel signal. Plant Physiol. 2001;126:1205–13.

11. Vogel JT, Walter MH, Giavalisco P, Lytovchenko A, Kohlen W, Charnikhova T, et al. SlCCD7 controls strigolactone biosynthesis, shoot branching and mycorrhiza-induced apocarotenoid formation in tomato. Plant J. 2010;61:300–11.

12. Kohlen W, Charnikhova T, Lammers M, Pollina T, Tóth P, Haider I, et al. The tomato CAROTENOID CLEAVAGE DIOXYGENASE8 (SlCCD8) regulates rhizosphere signaling, plant architecture and affects reproductive development through strigolactone biosynthesis. New Phytol. 2012;196:535–47.

13. Visentin I, Ferigolo LF, Russo G, Korwin Krukowski P, Capezzali C, Tarkowská D, et al. Strigolactones promote flowering by inducing the miR319-LA-SFT module in tomato. Proc Natl Acad Sci USA. 2024;121:e2316371121.

14. Sorefan K, Booker J, Haurogné K, Goussot M, Bainbridge K, Foo E, et al. MAX4 and RMS1 are orthologous dioxygenase-like genes that regulate shoot branching in Arabidopsis and pea. Genes Dev. 2003;17:1469–74.

15. Booker J, Sieberer T, Wright W, Williamson L, Willett B, Stirnberg P, et al. MAX1 encodes a cytochrome P450 family member that acts downstream of MAX3/4 to produce a carotenoid-derived branch-inhibiting hormone. Dev Cell. 2005;8:443–9.

16. Lin H, Wang R, Qian Q, Yan M, Meng X, Fu Z, et al. DWARF27, an iron-containing protein required for the biosynthesis of strigolactones, regulates rice tiller bud outgrowth. Plant Cell. 2009;21:1512–25.

17. Booker J, Auldridge M, Wills S, McCarty D, Klee H, Leyser O. MAX3/CCD7 is a carotenoid cleavage dioxygenase required for the synthesis of a novel plant signaling molecule. Curr Biol. 2004;14:1232–8.

18. Abe S, Sado A, Tanaka K, Kisugi T, Asami K, Ota S, et al. Carlactone is converted to carlactonoic acid by MAX1 in Arabidopsis and its methyl ester can directly interact with AtD14 in vitro. Proc Natl Acad Sci USA. 2014;111:18084–9.

19. Brewer PB, Yoneyama K, Filardo F, Meyers E, Scaffidi A, Frickey T, et al. LATERAL BRANCHING OXIDOREDUCTASE acts in the final stages of strigolactone biosynthesis in Arabidopsis. Proc Natl Acad Sci USA. 2016;113:6301–6.

20. Mashiguchi K, Seto Y, Onozuka Y, Suzuki S, Takemoto K, Wang Y, et al. A carlactonoic acid methyltransferase that contributes to the inhibition of shoot branching in Arabidopsis. Proc Natl Acad Sci USA. 2022;119:e2111565119.

21. Zhou A, Kane A, Wu S, Wang K, Santiago M, Ishiguro Y, et al. Evolution of interorganismal strigolactone biosynthesis in seed plants. Science. 2025;387:eadp0779.

22. Stirnberg P, Furner IJ, Ottoline Leyser HM. MAX2 participates in an SCF complex which acts locally at the node to suppress shoot branching. Plant J. 2007;50:80–94.

23. Arite T, Umehara M, Ishikawa S, Hanada A, Maekawa M, Yamaguchi S, et al. d14, a strigolactone-insensitive mutant of rice, shows an accelerated outgrowth of tillers. Plant Cell Physiol. 2009;50:1416–24.

24. Yao R, Ming Z, Yan L, Li S, Wang F, Ma S, et al. DWARF14 is a non-canonical hormone receptor for strigolactone. Nature. 2016;536:469–73.

25. Shabek N, Ticchiarelli F, Mao H, Hinds TR, Leyser O, Zheng N. Structural plasticity of D3-D14 ubiquitin ligase in strigolactone signalling. Nature. 2018;563:652–6.

26. Zhou F, Lin Q, Zhu L, Ren Y, Zhou K, Shabek N, et al. D14-SCF(D3)-dependent degradation of D53 regulates strigolactone signalling. Nature. 2013;504:406–10.

27. Soundappan I, Bennett T, Morffy N, Liang Y, Stanga JP, Abbas A, et al. SMAX1-LIKE/D53 Family Members Enable Distinct MAX2-Dependent Responses to Strigolactones and Karrikins in Arabidopsis. Plant Cell. 2015;27:3143–59.

28. Wang L, Wang B, Jiang L, Liu X, Li X, Lu Z, et al. Strigolactone Signaling in Arabidopsis Regulates Shoot Development by Targeting D53-Like SMXL Repressor Proteins for Ubiquitination and Degradation. Plant Cell. 2015;27:3128–42.

29. Tal L, Palayam M, Ron M, Young A, Britt A, Shabek N. A conformational switch in the SCF-D3/MAX2 ubiquitin ligase facilitates strigolactone signalling. Nat Plants. 2022;8:561–73.

30. Hu Q, Liu H, He Y, Hao Y, Yan J, Liu S, et al. Regulatory mechanisms of strigolactone perception in rice. Cell. 2024;187:7551–7567.e17.

31. Wang L, Wang B, Yu H, Guo H, Lin T, Kou L, et al. Transcriptional regulation of strigolactone signalling in Arabidopsis. Nature. 2020;583:277–81.

32. López-Ráez JA, Charnikhova T, Gómez-Roldán V, Matusova R, Kohlen W, De Vos R, et al. Tomato strigolactones are derived from carotenoids and their biosynthesis is promoted by phosphate starvation. New Phytol. 2008;178:863–74.

33. Umehara M, Hanada A, Yoshida S, Akiyama K, Arite T, Takeda-Kamiya N, et al. Inhibition of shoot branching by new terpenoid plant hormones. Nature. 2008;455:195–200.

34. Mayzlish-Gati E, De-Cuyper C, Goormachtig S, Beeckman T, Vuylsteke M, Brewer PB, et al. Strigolactones are involved in root response to low phosphate conditions in Arabidopsis. Plant Physiol. 2012;160:1329–41.

35. Gamir J, Torres-Vera R, Rial C, Berrio E, de Souza Campos PM, Varela RM, et al. Exogenous strigolactones impact metabolic profiles and phosphate starvation signalling in roots. Plant Cell Environ. 2020;43:1655–68.

36. Umehara M, Hanada A, Magome H, Takeda-Kamiya N, Yamaguchi S. Contribution of strigolactones to the inhibition of tiller bud outgrowth under phosphate deficiency in rice. Plant Cell Physiol. 2010;51:1118–26.

37. Xu X, Jibran R, Wang Y, Dong L, Flokova K, Esfandiari A, et al. Strigolactones regulate sepal senescence in Arabidopsis. J Exp Bot. 2021;72:5462–77.

38. Jing Y, Yang Z, Yang Z, Bai W, Yang R, Zhang Y, et al. Sequential activation of strigolactone and salicylate biosynthesis promotes leaf senescence. New Phytol. 2024;242:2524– 40.

39. Karniel U, Koch A, Bar Nun N, Zamir D, Hirschberg J. Tomato mutants reveal root and shoot strigolactone involvement in branching and broomrape resistance. Plants. 2024;13.

40. McCabe MS, Garratt LC, Schepers F, Jordi WJRM, Stoopen GM, Davelaar E, et al. Effects of PSAG12-*IPT* Gene Expression on Development and Senescence in Transgenic Lettuce. Plant Physiol. 2001;127:505–16.

41. Li M, Li X, Du J, Li W, He R, Lin Y, et al. Effects of exogenous application of the strigolactone GR24 on quality and flavor components during postharvest storage of celery. Postharvest Biol Technol. 2024;212:112900.

42. Scaffidi A, Waters MT, Sun YK, Skelton BW, Dixon KW, Ghisalberti EL, et al. Strigolactone Hormones and Their Stereoisomers Signal through Two Related Receptor Proteins to Induce Different Physiological Responses in Arabidopsis. Plant Physiol. 2014;165:1221–32.

43. Zhao Y, Zhang Y, Li S, Tan S, Cao J, Wang H-L, et al. Leaf Senescence Database v5.0: A Comprehensive Repository for Facilitating Plant Senescence Research. J Mol Biol. 2024;436:168530.

44. Wu J, Li Y, Yin H, Zhao L, Xu C, Fu X, et al. Plant heat shock protein Hsp90 enhances stress resistance through integrating protein quality control, chloroplast protection, hormone signal network, and immune defense. J Exp Bot. 2026;77:910–31.

45. Panaretou B, Siligardi G, Meyer P, Maloney A, Sullivan JK, Singh S, et al. Activation of the ATPase activity of hsp90 by the stress-regulated cochaperone aha1. Mol Cell. 2002;10:1307–18.

46. Belisle CE, Sargent SA, Brecht JK, Sandoya GV, Sims CA. Accelerated Shelf-life Testing to Predict Quality Loss in Romaine-type Lettuce. Horttechnology. 2021;31:490–9.

47. Inskeep WP, Bloom PR. Extinction coefficients of chlorophyll a and B in n,n-dimethylformamide and 80% acetone. Plant Physiol. 1985;77:483–5.

48. Van Bel M, Silvestri F, Weitz EM, Kreft L, Botzki A, Coppens F, et al. PLAZA 5.0: extending the scope and power of comparative and functional genomics in plants. Nucleic Acids Res. 2022;50:D1468–74.

49. Letunic I, Bork P. Interactive Tree of Life (iTOL) v6: recent updates to the phylogenetic tree display and annotation tool. Nucleic Acids Res. 2024;52:W78–82.

50. Engler C, Youles M, Gruetzner R, Ehnert T-M, Werner S, Jones JDG, et al. A golden gate modular cloning toolbox for plants. ACS Synth Biol. 2014;3:839–43.

51. Martin M. Cutadapt removes adapter sequences from high-throughput sequencing reads. EMBnet j. 2011;17:10.

52. Reyes-Chin-Wo S, Wang Z, Yang X, Kozik A, Arikit S, Song C, et al. Genome assembly with *in vitro* proximity ligation data and whole-genome triplication in lettuce. Nat Commun. 2017;8:14953.

53. Dobin A, Davis CA, Schlesinger F, Drenkow J, Zaleski C, Jha S, et al. STAR: ultrafast universal RNA-seq aligner. Bioinformatics. 2013;29:15–21.

54. Smith T, Heger A, Sudbery I. UMI-tools: modeling sequencing errors in Unique Molecular Identifiers to improve quantification accuracy. Genome Res. 2017;27:491–9.

55. Liao Y, Smyth GK, Shi W. featureCounts: an efficient general purpose program for assigning sequence reads to genomic features. Bioinformatics. 2014;30:923–30.

56. Love MI, Huber W, Anders S. Moderated estimation of fold change and dispersion for RNA-seq data with DESeq2. Genome Biol. 2014;15:550.

57. Ritter A, Iñigo S, Fernández-Calvo P, Heyndrickx KS, Dhondt S, Shi H, et al. The transcriptional repressor complex FRS7-FRS12 regulates flowering time and growth in Arabidopsis. Nat Commun. 2017;8:15235.

58. Waibel F, Filipowicz W. U6 snRNA genes of Arabidopsis are transcribed by RNA polymerase III but contain the same two upstream promoter elements as RNA polymerase II-transcribed U-snRNA genes. Nucleic Acids Res. 1990;18:3451–8.

59. Hu X, Meng X, Liu Q, Li J, Wang K. Increasing the efficiency of CRISPR-Cas9-VQR precise genome editing in rice. Plant Biotechnol J. 2018;16:292–7.

60. Concordet J-P, Haeussler M. CRISPOR: intuitive guide selection for CRISPR/Cas9 genome editing experiments and screens. Nucleic Acids Res. 2018;46:W242–5.

61. Moreno-Mateos MA, Vejnar CE, Beaudoin J-D, Fernandez JP, Mis EK, Khokha MK, et al. CRISPRscan: designing highly efficient sgRNAs for CRISPR-Cas9 targeting *in vivo*. Nat Methods. 2015;12:982–8.

62. Michelmore R, Marsh E, Seely S, Landry B. Transformation of lettuce (Lactuca sativa) mediated by Agrobacterium tumefaciens. Plant Cell Rep. 1987;6:439–42.

