## Supplemental Figure 1 and 2 for "Loss of MAX1 redirects, rather than delays, the leaf senescence program in lettuce"

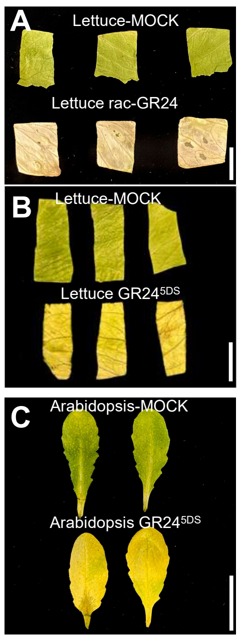


**D**

| *Functionality* | *Name of gene* | *Arabidopsis thaliana* | *Lactuca sativa* |
|---|---|---|---|
| Biosynthesis | D27 | AT1G03055 | Lsat_1_v5_gn_8_94460<br>Lsat_1_v5_gn_9_7981 |
| | MAX3 | AT2G44990 | Lsat_1_v5_gn_3_130581<br>Lsat_1_v5_gn_3_130781<br>Lsat_1_v5_gn_3_129760 |
| | MAX4 | AT4G32810 | Lsat_1_v5_gn_1_46661<br>Lsat_1_v5_gn_6_23181 |
| | MAX1 | AT2G16170 | Lsat_1_v5_gn_6_14580 |
| | LBO | AT3G21420 | Lsat_1_v5_gn_6_14981 |
| | CLAMT | AT4G36470 | Lsat_1_v5_gn_3_75880 |
| | CYP722A1 | AT1G19630 | Lsat_1_v5_gn_3_77380<br>Lsat_1_v5_gn_3_56180 |
| Signaling | D14 | AT3G03990 | Lsat_1_v5_gn_1_24401 |
| | MAX2 | AT2G42620 | Lsat_1_v5_gn_8_141221 |
| | SMXL6 | AT1G07200 | Lsat_1_v5_gn_5_171201 |
| | SMXL7 | AT2G29970 | |
| | SMXL8 | AT2G240130 | Lsat_1_v5_gn_2_123560 |

**E**

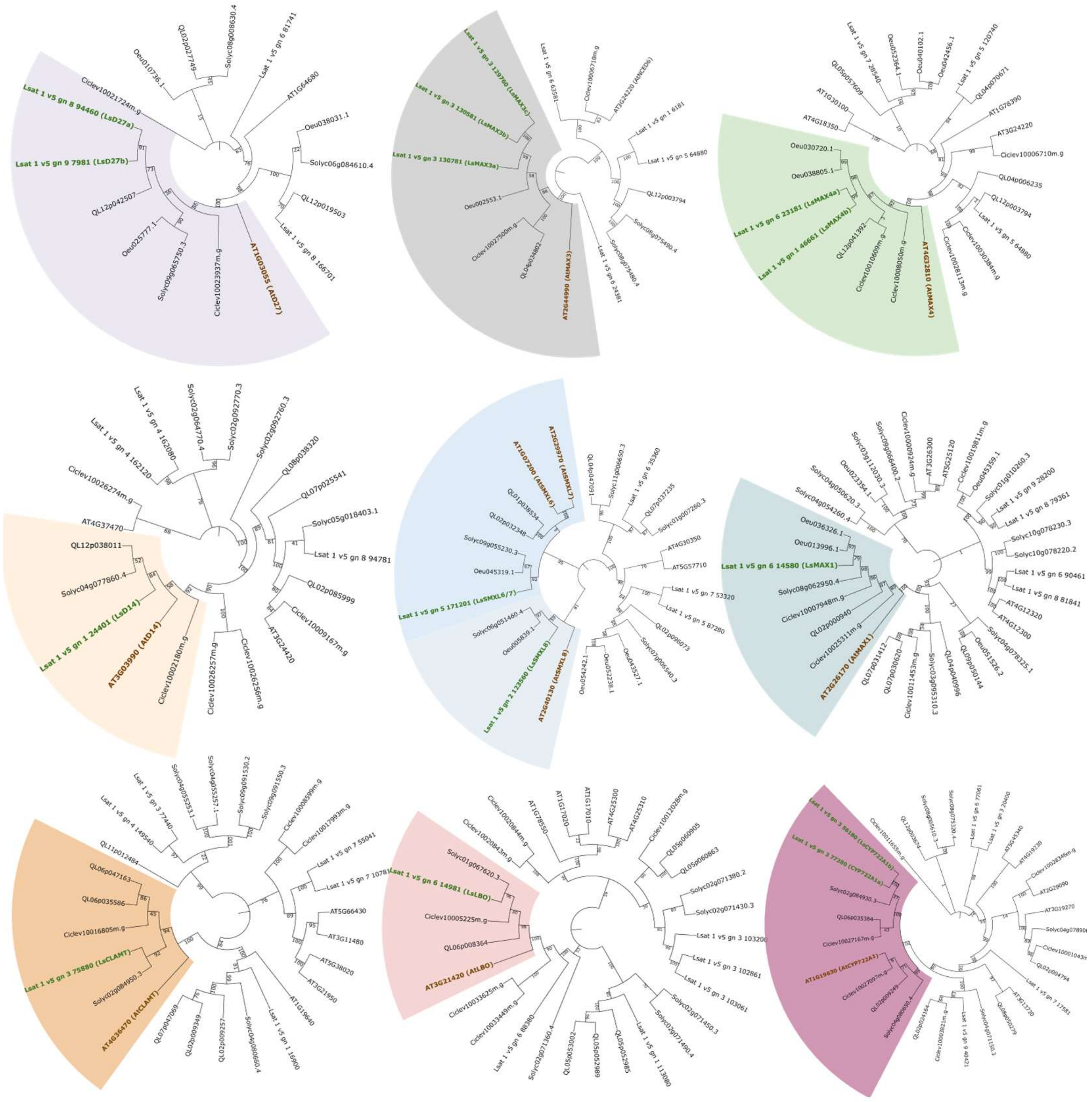

**Supplementary Figure 1**. **SL pathway conservation and senescence activity in lettuce**. **A.** Lettuce leaf discs, treated with either acetone or 10 μM racemic-GR24 dissolved in acetone, following dark incubation. scale bar = 2 cm. **B.** Lettuce leaf discs, treated with either acetone or 10 μM GR24$^{5DS}$ dissolved in acetone, following dark incubation. scale bar = 2 cm. **C.** Arabisopsis leaf , treated with either acetone or 10 μM GR24$^{5DS}$ dissolved in acetone, following dark incubation. scale bar = 2 cm **D.** Lettuce homologs of strigolactone signaling and biosynthesis pathways **E.** Phylogenetic tree constructed from protein sequences from *Lactuca sativa, Arabidopsis thaliana, Solanum lycopersicum, Quercus lobata, Olea europaea, and Citrus clementina*. Arabidopsis thaliana SL genes are indicated in brown, and the Lactuca sativa homolog is indicated in green.

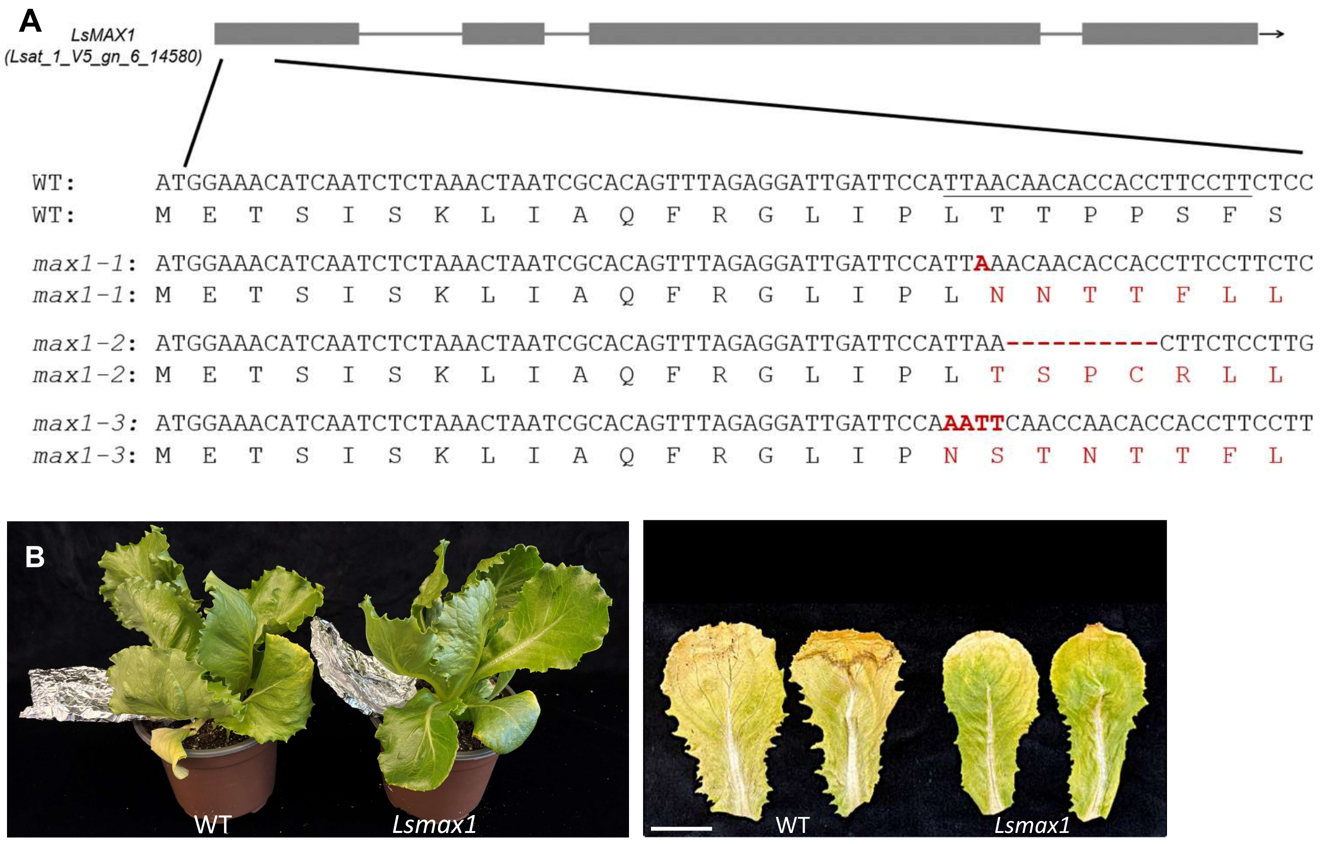


**Supplementary Figure 2**. **CRISPR/Cas9-generated *Lsmax1* alleles with delayed senescence phenotype. A**. Schematic representation of the *LsMAX1* gene structure with exon-intron organization (top). Partial nucleotide and corresponding amino acid sequences of WT and *Lsmax1* CRISPR/CAS9-generated mutant alleles (*Lsmax1-1*, *Lsmax1-2*, and *Lsmax1-3*) are shown below. Mutations relative to WT are indicated in red; dashes denote deletions. Predicted amino acid changes resulting from each mutation are shown beneath the nucleotide sequences. **B**. WT and *Lsmax1* plants with the fourth leaf covered with aluminum foil for 14 days. Left: whole-plant view showing the covered leaf. Right: close-up of the treated leaf after 14 days of coverage. Scale bar = 3 cm.
